# Imputation aware tag SNP selection to improve power for multi-ethnic association studies

**DOI:** 10.1101/105551

**Authors:** Genevieve L. Wojcik, Christian Fuchsberger, Daniel Taliun, Ryan Welch, Alicia R Martin, Suyash Shringarpure, Christopher S. Carlson, Goncalo Abecasis, Hyun Min Kang, Michael Boehnke, Carlos D. Bustamante, Christopher R. Gignoux, Eimear E. Kenny

**Affiliations:** Department of Genetics, Stanford University School of Medicine, Stanford, CA, USA; Department of Biostatistics and Center for Statistical Genetics, School of Public Health, University of Michigan, Ann Arbor, MI, USA; Fred Hutchinson Cancer Center, University of Washington, Seattle, WA, USA; Department of Biomedical Data Science, Stanford University School of Medicine, Stanford, CA, USA; Department of Genetics and Genomic Sciences, Icahn School of Medicine at Mount Sinai, New York, NY, USA; The Charles Bronfman Institute for Personalized Medicine, Icahn School of Medicine at Mount Sinai, New York, NY, USA; The Icahn Institute of Multiscale Biology and Genomics, Icahn School of Medicine at Mount Sinai, New York, NY, USA; The Center for Statistical Genetics, Icahn School of Medicine at Mount Sinai, New York, NY, USA

## Abstract

The emergence of very large cohorts in genomic research has facilitated a focus on genotype-imputation strategies to power rare variant association. Consequently, a new generation of genotyping arrays are being developed designed with tag single nucleotide polymorphisms (SNPs) to improve rare variant imputation. Selection of these tag SNPs poses several challenges as rare variants tend to be continentally-or even population-specific and reflect fine-scale linkage disequilibrium (LD) structure impacted by recent demographic events. To explore the landscape of tag-able variation and guide design considerations for large-cohort and biobank arrays, we developed a novel pipeline to select tag SNPs using the 26 population reference panel from Phase of the 1000 Genomes Project. We evaluate our approach using leave-one-out internal validation via standard imputation methods that allows the direct comparison of tag SNP performance by estimating the correlation of the imputed and real genotypes for each iteration of potential array sites. We show how this approach allows for an assessment of array design and performance that can take advantage of the development of deeper and more diverse sequenced reference panels. We quantify the impact of demography on tag SNP performance across populations and provide population-specific guidelines for tag SNP selection. We also examine array design strategies that target single populations versus multi-ethnic cohorts, and demonstrate a boost in performance for the latter can be obtained by prioritizing tag SNPs that contribute information across multiple populations simultaneously. Finally, we demonstrate the utility of improved array design to provide meaningful improvements in power, particularly in trans-ethnic studies. The unified framework presented will enable investigators to make informed decisions for the design of new arrays, and help empower the next phase of rare variant association for global health.

## Introduction

The vast majority of human genomic variation is rare ^1^, and an appreciable fraction of rare variants are likely to be functionally consequential. ^2^ The gold standard approach to assay rare variation is via sequencing. So far, large-scale sequencing studies have had some, but limited, success for discovery of rare variant associations^3–6^. There is a new appreciation that studies of hundreds of thousands or millions of individuals will be needed to drive well-powered discovery efforts. ^7,8^ Currently, genome sequencing on this scale is prohibitively expensive and computationally burdensome. In contrast, genome-wide genotyping arrays are inexpensive, with far less bioinformatic overhead compared to sequencing. The past decade of genomic research has seen the development of myriad commercial high-throughput genotyping arrays. ^9,10^ While initially designed to capture common variants ^11^, in recent years arrays have been leveraged to capture variation at the rare end of the frequency spectrum. One strategy is to ascertain rare variants directly on arrays, which is restricted to a very narrow subset of the rare variant spectrum due to array size limits.^12–14^ Another strategy is to leverage the haplotype structure determined by common variants on the array, which form a ‘scaffold’, for accurate inference of un-genotyped variation through multi-marker imputation into sequenced reference panels of whole genomes. The strategy of genotyping, followed by imputation, has the potential to recover rare untyped variants in very large cohorts of arrayed samples at no additional experimental cost. ^15,16^ This model bridging genotyping and imputation has prompted efforts to build deep reference sequence databases and a renewed interest in methods for improving genome-wide scaffold design. ^6,17,18^

Genotype array scaffolds have been designed historically using algorithms that select tagging single nucleotide polymorphisms (tag SNPs) that are in linkage disequilibrium (LD) with a maximal number of other SNPs. Tag SNP algorithms are optimized to maximize this score, typically described as pairwise coverage. However, imputation tools increasingly incorporate sophisticated haplotype information to impute unobserved variants.^19–21^ Consequently, it is not clear that tag SNPs that maximize pairwise coverage will be tag SNP's that provide, in aggregate, the best GWAS scaffold for accurate imputation. ^22^ Further, most tag SNP selection algorithms use LD architecture in a single population, ^23,24^, while we know LD patterns can vary extensively between populations.^17^ Historically, many commercial arrays were designed by selecting tag SNPs from European populations, although arrays targeting some other populations have recently entered the market. ^9,10^ The number of SNPs tagged by a tag SNP can vary appreciably between populations due to demographic forces of migration, population expansion, and genetic drift. This may diminish GWAS scaffold performance in populations other than those in which the tag SNPs were selected, which in turn, can lead to reduced power for imputation-based association. This is a particularly pernicious problem in populations for which no targeted commercial array is available, in studies with multi-ethnic populations, and for accurate estimation of the transferability of genetic risk across populations.

As association studies grow larger and increasingly diverse, there is a need to reassess design criteria for GWAS scaffolds and arrays. ^25,26^ On the one hand, tag SNPs that tag lower frequency variants are likely to be on the lower end of the site frequency spectrum and, consequentially, more geospatially restricted.^27–30^ On the other hand, as studies grow very large, cohort heterogeneity is likely to increase substantially for both discovery and replication populations. ^31,32^ Given finite GWAS scaffold density, examining the trade-off between lowering the frequency threshold for accurate imputation and extending utility to multiple populations will become important. ^33,34^ In this manuscript, we describe a framework for developing well-powered tag SNP selection leveraging thousands of whole genomes from diverse populations for balanced cross-population coverage. In our study, genomic coverage is evaluated based on genome-wide imputation accuracy as measured by mean imputed r^2^ at untyped sites, rather than pairwise linkage disequilibrium. Moving beyond pairwise metrics allows us to account for haplotype diversity across the genome and demonstrates population-specific biases from pairwise estimates. Assessing accuracy using leave-one-out cross-validation yields a real-world estimate of genomic coverage. We examine the effect of allele frequency, correlation thresholds, and population diversity on the selection of tag SNP and on the landscape of tag-able variation. This work demonstrates that, while there may be limits given current reference panels, improving GWAS scaffold design is an underused means to increase power in association studies.

## Results

### Selecting tag SNPs from a single population results in suboptimal tagging performance across populations impacted by different patterns of demography

First we designed an experiment to assess imputation accuracy performance comparing tag SNP selection from different populations. This experiment mimics the current design of many commercial arrays, in which tag SNPs were selected to capture the primarily variation in a single population or a closely related group of populations. We built a pipeline using the 26 population reference panel from Phase 3 of the 1000 Genomes Project and the Tagit algorithm for tag SNP selection.^23^
**(Supplementary Table 1)** Individuals were split into mutually exclusive “super populations.” These included the Admixed American (AMR), East Asian (EAS), European (EUR), and South Asian (SAS) populations as described in Auton et al.^17^ In addition, we divided the African super population into two groups: four populations from Africa (AFR) and two populations of African 19,21,35-38 descent in the Americas (AAC) (see **Methods**). Initially, to mimic the design of many arrays, tag SNPs were only selected from a single super population. We assumed a genome-wide allocation of 500,000 tag SNPs, however analyses for a single population tagging strategy were only conducted on chromosome 9 with the allocation of 21,107 sites proportional to the physical distance of chromosome 9 compared to all chromosomes combined. Potential tags were required to have a minor allele frequency (MAF) ≥ 1% and be in pairwise LD with the tagged target site with a r^2^ ≥ 0.5.

The current generation of phase-based imputation algorithms (BEAGLE, IMPUTE2, Minimac3) leverage local haplotype information and sequenced reference panels to improve accuracy of variant inference compared to tag SNP approaches. ^19–,3538^ Therefore, optimal array design depends not only on tag SNP selection, but also on empirical evaluation of imputation performance. For each of the population-specific GWAS scaffolds, imputation accuracy was assessed in all six super populations by MAF bins (common, MAF = 0.05-0.5; low frequency, MAF = 0.01-0.05; and rare, MAF < 0.01) by comparing the imputed dosages to the real genotypes through leave-one-out internal validation. (see Methods)

Consistently across all super populations, the population from which the tags were ascertained had the highest imputation accuracy in the common bin. **(Supplementary Figure 1)** Trends in imputation accuracy follow known patterns of demography. For example, if the tags were ascertained in European populations, imputation accuracy was best in Europeans (EUR), followed by out-of-Africa populations (AMR, SAS, EAS), and worst in African ancestry populations (AFR, AAC). **(Figure 1)** If the tags were ascertained in African populations, the inverse was observed. (**Supplementary Figure 1**) As expected, the same trend of reduced imputation accuracy in non-ascertained populations was exacerbated in the low frequency bin. Imputation of low frequency variants in East Asian populations (EAS) was consistently most challenging; even when tag SNPs were selected from EAS, accuracy of low frequency imputation was the same or better in other populations. This can be explained by evidence of a recent tight bottleneck followed by rapid population grown in EAS, resulting in a large proportion of rare variants that are difficult to tag due to lower LD, especially with a limited scaffold of 500,000 sites. ^27^ In contrast, the imputation performance of tag SNPs ascertained in AFR, AMR, and AAC populations is the same or better compared to the performance in out-of-Africa populations. This is likely due to increased allelic heterogeneity in African ancestry populations, which results in greater haplotypic diversity and a higher chance that a rare variant is well tagged by a haplotype for imputation.^17^ The imputation accuracy of AMR higher in the rare frequency bin (MAF 0.5-1%), independent of the ascertainment population, is likely due to longer haplotypes resulting from recent admixture, allowing the rare variation to be captured accurately given the limited allocation.^39^ Importantly, in each case we observe a notable drop-off in performance across most of the frequency spectrum when examiningimputation coverage in populations diverging from the one used for tag SNP selection **(Supplementary Figure 1)**.

**Figure 1:**
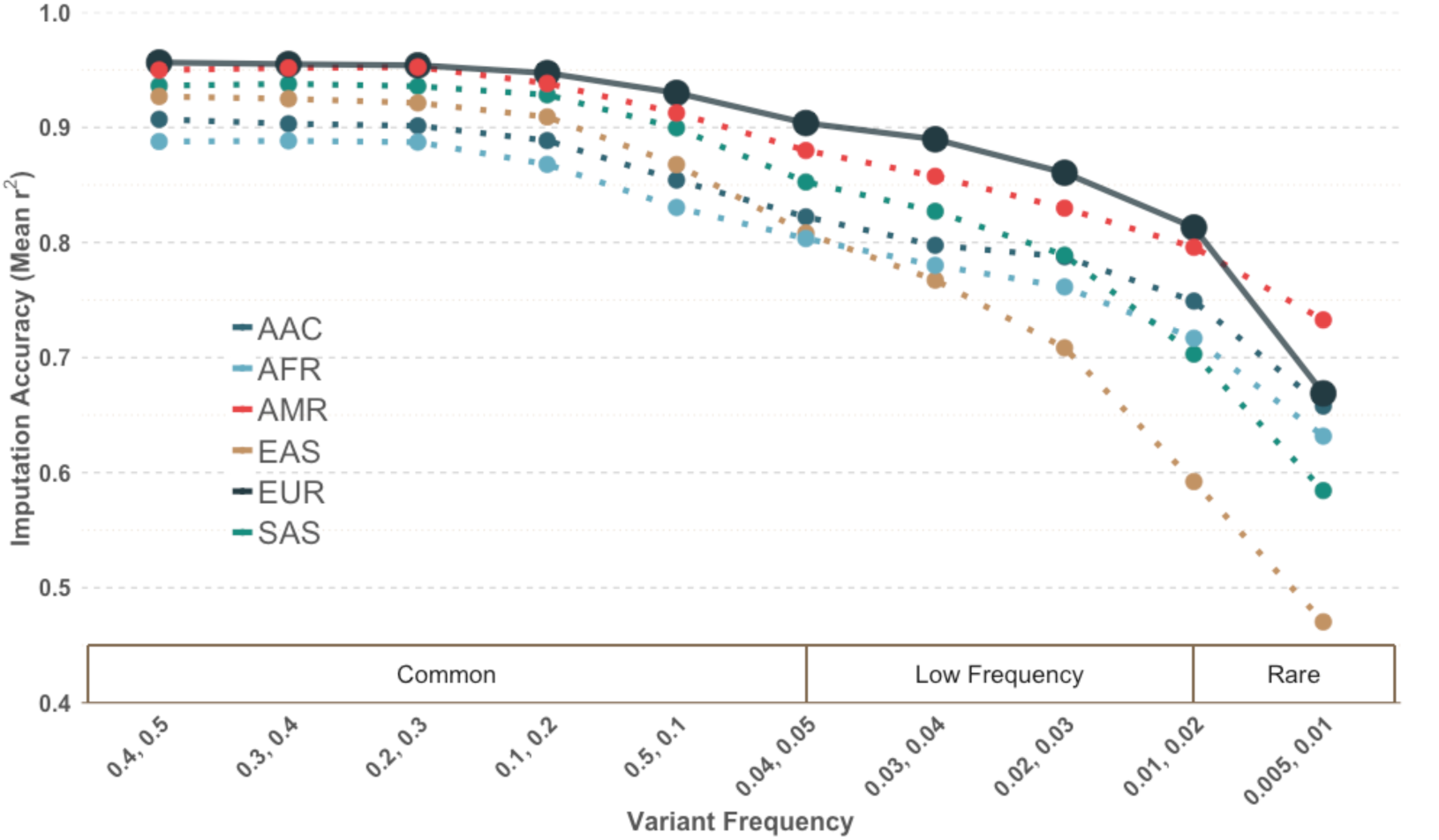
Imputation Accuracy by super population of tags selected in European populations for a scaffold assuming 500,000 genome-wide variants. Tags were required to have a MAF≥1% and r^2^≥0.5 with target sites. This trend is observed across all super populations (*Supplemental Figure 1*).

## Comparing single versus cross population tag SNP selection strategies

The first criterion of developing a useful genotyping platform is whether or not the variants assayed segregate in the population of interest and contribute tagging ability by being in LD (high r^2^) with untagged sites. For example, using <u>Illumina’s OmniExpress platform</u> within the 1000 Genomes Project data, over 99.7% of the sites will be polymorphic (MAF>0.5%) in the overall dataset. However, when we stratify by super population, each group has a differential loss. While AFR loses <1% of sites for having a MAF<0.5%, EUR and EAS lose 4.4% and 9.2% of variants, respectively. This will lead to a loss of statistical power dependent on ancestry and could limit analyses. This is quantified as “informativeness”, or the ability of a tag SNP to both segregate in the population and provide LD information (r^2^>0.5 with at least one untagged site). When working in multiple populations, it is essential to have balanced representation of variation across all groups.

To explore different approaches for GWAS scaffold design we compared three strategies for selecting tag SNPs; single population tag SNP ascertainment, in which all tags are selected from a single population; a ‘naïve’ approach, in which all populations are combined and tags are selected based on composite statistics derived from this multi-population pool; and a ‘cross-population prioritization’ approach, in which tags are prioritized if they are both informative in multiple populations and by the number of unique sites targeted across all groups (see **Methods** and **Supplementary Figure 2**). We generated lists of tags per method assuming a total genome-wide allocation of 500,000 sites and minimum thresholds of r^2^>0.5 and minor allele frequency (MAF) ≥ 1%. Using these parameters, an exhaustive set of tag SNPs were selected using the naïve approach with tags ranked by the absolute number of sites tagged across the 6 super populations, regardless of how many super populations had LD between tags and targets. We then re-ranked them using the cross-population prioritization approach (**Supplementary Figure 2**).

To compare the three approaches, we tallied the number of informative tags per population for each method to investigate the added value of tags contributing information in multiple populations. **(Figure 2)** This was done for all 22 autosomes. As per the design, all the single-population tags were informative within the super population from which tag SNPs were selected. Comparing the naïve and cross-population approaches that selected tag SNPs across all populations, the cross-population prioritization approach increased the number of informative tag SNPs in all populations relative to the naïve approach. In the naïve approach, we observed that the majority of tag SNPs were selected from the AFR population, followed by AAC, due to African-descent populations having more polymorphic sites across the genome with lower linkage disequilibrium. ^17,40^ Whereas in the cross-population prioritization approach, variation specific to a single population is down-weighted leading to more balanced representation between all 6 super populations. By leveraging cross-population information the largest boost in the proportion of tag SNPs contributing linkage disequilibrium information compared to the naive approach was observed in non-African descent populations (10.5%, 28.6%, 25.9%, and 28.7% in AMR, EAS, EUR and SAS, respectively). Even the African descent populations (AFR and AAC), which dominate the naïve approach, have a higher proportion of tags in linkage disequilibrium with target sites with the cross-population prioritization approach (a 2.2% and 1.0% boost for AAC and AFR, respectively).

**Figure 2:**
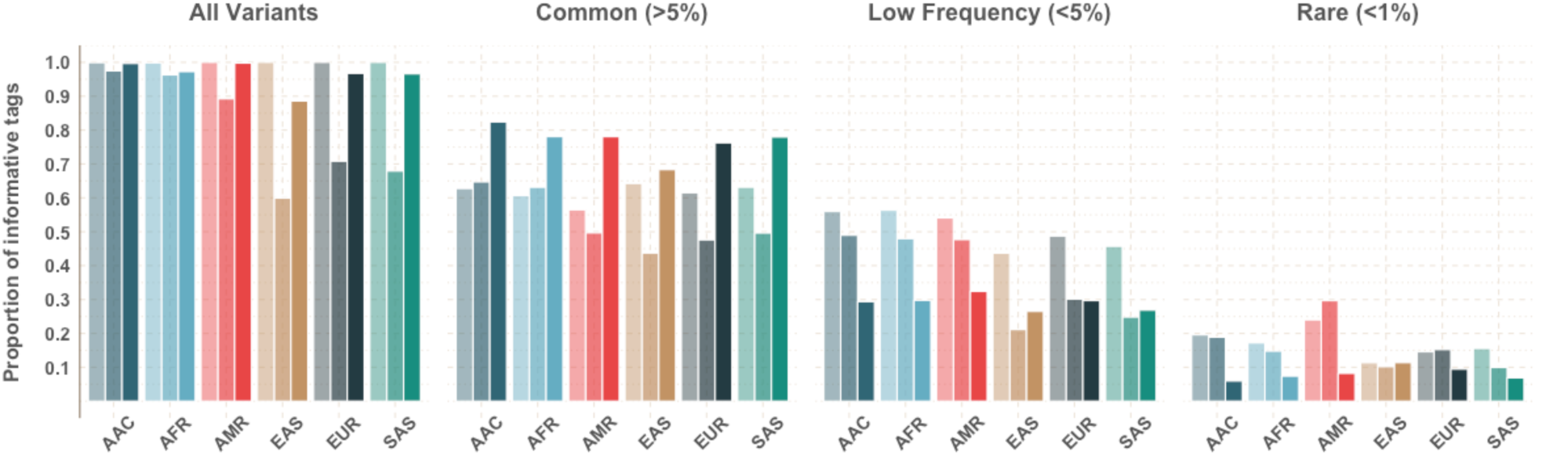
Proportion of tags that are informative by population with the three methods. (Left, lightest)tags selected from only a single population, (Center) tags selected by pooling all populations agnostically, and (Right) tags selected with the cross-population prioritization approach. Tag SNPs were informative if they were in linkage disequilibrium (r^2^>0.5) with at least one untagged site.

To assess performance across the frequency spectrum we also stratified our accuracy estimates by super population-specific MAF into common, low frequency, and rare bins, as previously described. We observed that the cross-prioritization approach results in a larger proportion of tags being informative compared to both the single-population and naïve for common tag SNPs (MAF>0.05) in all super populations. This is likely because the cross-prioritization approach prioritizes potential tag SNPs that provide LD information across multiple populations, therefore prioritizing common variants tagging common variation. However, by limiting tag SNP selection to these common variants only, the proportion of tags that provide LD information for low frequency variants is decreased compared to the single population approach, which had the highest proportion of informative tag SNPs in low and rare frequency in the target population. For example, when tags were ascertained using only AAC LD information, 19.5% of the 500,000 SNP scaffold were informative for rare variation (MAF<1%) and 62.8% for common variation MAF>5%) within AAC populations. When the cross-population approach was used, ensuring the prioritization of common variation, the proportion of tag SNPs informative for rare variation dropped to 6% while the proportion informative for common variation jumped up to 82.4%. This is consistent with low frequency and rare variants being population-specific, therefore not tagged by cosmopolitan common variation present in multiple populations. A notable exception is that the naïve approach contributes the most LD information for rare variants in the AMR super population. This is consistent with our previous findings showing highest imputation accuracy in the rare variation within AMR, even when the population from which tag SNPs were ascertained was different. The AMR on average exhibit longer haplotype lengths from the recently admixed populations in the Americas. ^17,39^ Because of the long haplotype tract lengths, more limited haplotypic diversity, and the limited allocation of tag SNPs, a naïve approach emphasizing the absolute number of unique sites up-weights variation that is informative for at least one of the ancestral components present in these populations.

### Cross population prioritization of tag SNPs increases imputation accuracy for all groups across frequency spectrum compared to naïve approach

Beyond being polymorphic and providing LD information across global populations, an efficient tag SNP scaffold must also optimize the LD structure beyond individual tag SNPs’ pairwise. The goal of tag SNP selection is to inform the unmeasured haplotypes, and therefore their performance must be evaluated as a collaborative unit. One way to assess this is through imputation accuracy. Hence, following the observation that cross-population prioritization selects a higher proportion of informative common tag SNPs for each population, even compared to the single population approach, we next assessed what impact this would have on imputation accuracy. We deployed the same leave-one-out internal cross validation approach as before using the 1000 Genomes Project populations (see **Methods**). We assumed a genome-wide scaffold of 500,000 sites and tags had to have a MAF>1% and r^2^>0.5 with tagged sites. Imputation accuracy was highest across all population-specific minor allele frequency bins when ascertaining in the target population in non-African non-admixed descent continental populations (EAS, EUR, and SAS). **(Supplementary Figure 3)** For the two African descent groups (AAC and AFR), the cross-population prioritization approach had the highest imputation accuracy across all sites. When stratified by MAF bins, the increase in informative tag SNPs for common variants with the cross population approach yielded higher imputation accuracy for common variation in all super populations. As previously seen, the population-specific nature of low frequency and rare variants led to decreased imputation accuracy in non-African descent populations for both the cross-population and naïve approach when compared to targeted single-population ascertainment. The cross-population prioritization approach had higher imputation accuracy than the naïve approach for all MAF bins.

As scaffold size can dramatically affect imputation accuracy^41^, we additionally examined allocations of 250,000, 500,000, 1,000,000, 1,500,000, and 2,000,000 genome-wide tags, which were all selected with r^2^>0.5 and MAF>0.01. The cross-population prioritization scheme performed better with higher imputation accuracy than the naïve method for all super populations across all minor allele frequency bins with tags selected. **(Figure 3)** The biggest improvement came with the smaller array sizes. The most marked improvement was found in EAS, which originally had the lowest imputation accuracy of the 6 super populations with the naive approach. Within EAS groups, the cross-population approach increased imputation accuracy overall by 9.8% (from 67.3% to 77.1%) for a tag scaffold of 250,000 sites. For a scaffold of 500,000 sites, an overall improve of 6.2% was observed (from 77.4% to 83.6%). Improvements were largely consistent with the increase of informative tag SNPs. **(Figure 2)** As with the naive prioritization approach SNPs were disproportionately informative within AFR and AAC, consistent with admixed ancestry reflected by reference panels. For the smaller sizes (250K), the greatest increase in performance incorporating cross-population information was found within common SNPs (MAF>5%). However, the larger sized scaffolds (1-2 million) showed the most improvement within the low frequency bins (MAF<5%).

**Figure 3:**
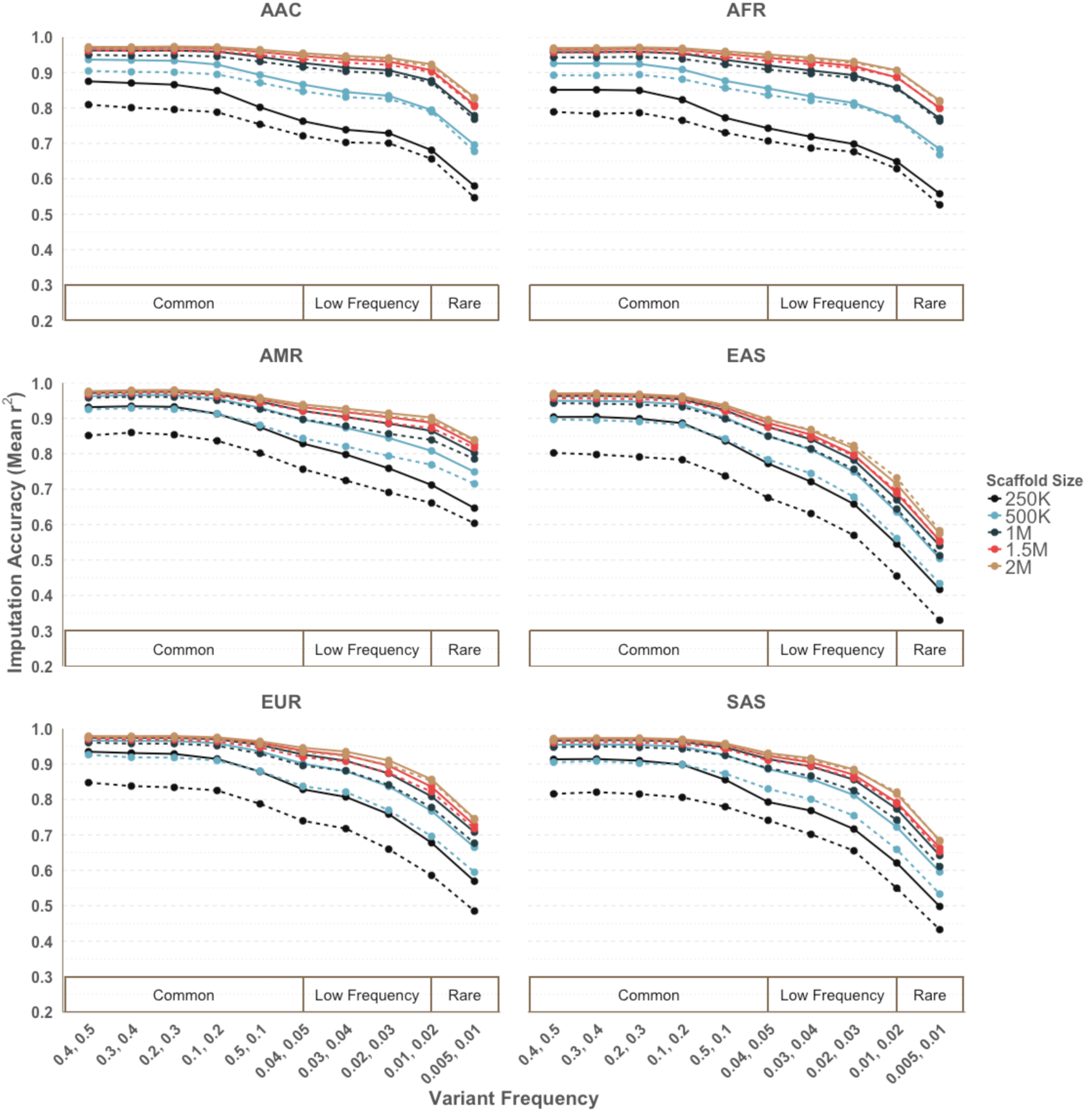
Increased imputation accuracy with cross-population prioritization (solid line) versus naïve approach (dashed line) for a minimum pairwise correlation threshold of r^2^>0.5 and MAF>1% across different scaffold sizes. Imputation accuracy was calculated separately within minor allele frequency bins for each super population.

### Imputation accuracy varies by local ancestry background in admixed individuals

We also assessed imputation ancestry stratified by local ancestry diplotype in the two admixed populations, the AAC and AMR, for a genome-wide allocation of 500,000 tag SNPs. First, using phased data, we inferred haploid tracts of African, European, and Native American local ancestry along the genomes of all individuals in the AMR and AAC populations (see **Methods**, ^17,42^). Then each variant was inferred to be on one of six ancestral diploid tracts; European-European (EUR-EUR), European-African (EUR-AFR), European-Native American (EUR-NAT), African-Native American (AFR-NAT), African-African (AFR-AFR) and Native American-Native American (NAT-NAT). In all local ancestry strata the cross-population prioritization yielded improved imputation accuracy when compared to the naïve approach. When looking at ASW population (Americans of African ancestry in South West US), performance was high overall with all diploid tracts having imputation accuracies of 92.8-96.8% for all sites with minor allele frequency above 1%. **(Supplementary Figure 4)** The lowest imputation accuracy was found in AFR-AFR tracts, especially at the lower end of the frequency spectrum. The highest imputation accuracy was found in EUR-EUR tracts (94% overall for ASW). In AMR populations, by contrast, the NAT-NAT tracts had the lowest performance of all. An example can be seen in the MXL population (Mexican Ancestry from Los Angeles), where the highest imputation accuracy was found in the AFR-EUR tracts (overall imputation accuracy of 90.1% for all SNPs with MAF>0.5%) and the lowest within NAT-NAT tracts (74.8% for all SNPS with MAF>0.5%). **(Supplementary Figure 4B)** These performances are reflective of the relative availability of reference data relevant to these specific ancestral components.

### Evaluating impact of r^2^ and MAF thresholds on tag SNP performance

Previous standards in scaffold design have considered minimum linkage disequilibrium (r^2^) and minor allele frequency (MAF) thresholds when prioritizing possible tag SNPs. However, the impact of these thresholds are often evaluated through pairwise coverage. We explored varying the minimum r^2^ threshold (0.2, 0.5, 0.8) and MAF (0.5%, 1%, 5%) to assess their impacts on imputation accuracy, as well as pairwise coverage, assuming a genome-wide allocation of one million tags. For common variants, a higher minimum r^2^ threshold (r^2^>0.8) resulted in slightly higher imputation accuracy. **(Figure 4A)** However, the sites in the low and rare bin demonstrate population-specific accuracy only. **(Supplementary Figure 5)** For AFR, SAS, and EAS, a less stringent threshold of r^2^>0.2 had the worst imputation accuracy across all frequency bins. Low frequency and rare variation had higher imputation accuracy for an r^2^ threshold of 0.5 compared to 0.8. Within AAC, AMR, and EUR, the low frequency variation had improved imputation accuracy with the lowest r^2^ threshold of 0.2. However, the imputation accuracy within this low threshold was notably compromised for common variants. This indicates that low frequency variation is better captured by weak correlation structure, but at a cost to common variation in these populations. Analyses performed with r^2^>0.5 had the best balance of performance across all frequency bins with the highest overall imputation accuracy in all super populations except for EAS. **(Supplementary Table 2)** Overall, there was very small differences in imputation accuracy between the different r^2^ thresholds. There were much larger differences in coverage, including both coverage evaluated with minimum r^2^ (LD) of 0.5 and 0.8. **(Figure 4A)** Additionally, the best “performance” using pairwise coverage was highly dependent on the definition of coverage. Specifically, if pairwise coverage wascalculated as the proportion of sites that are in LD with r^2^>0.5, then the best minimum r^2^ threshold in tag SNP selection will be 0.5. This holds true for r^2^>0.8 as well.

**Figure 4:**
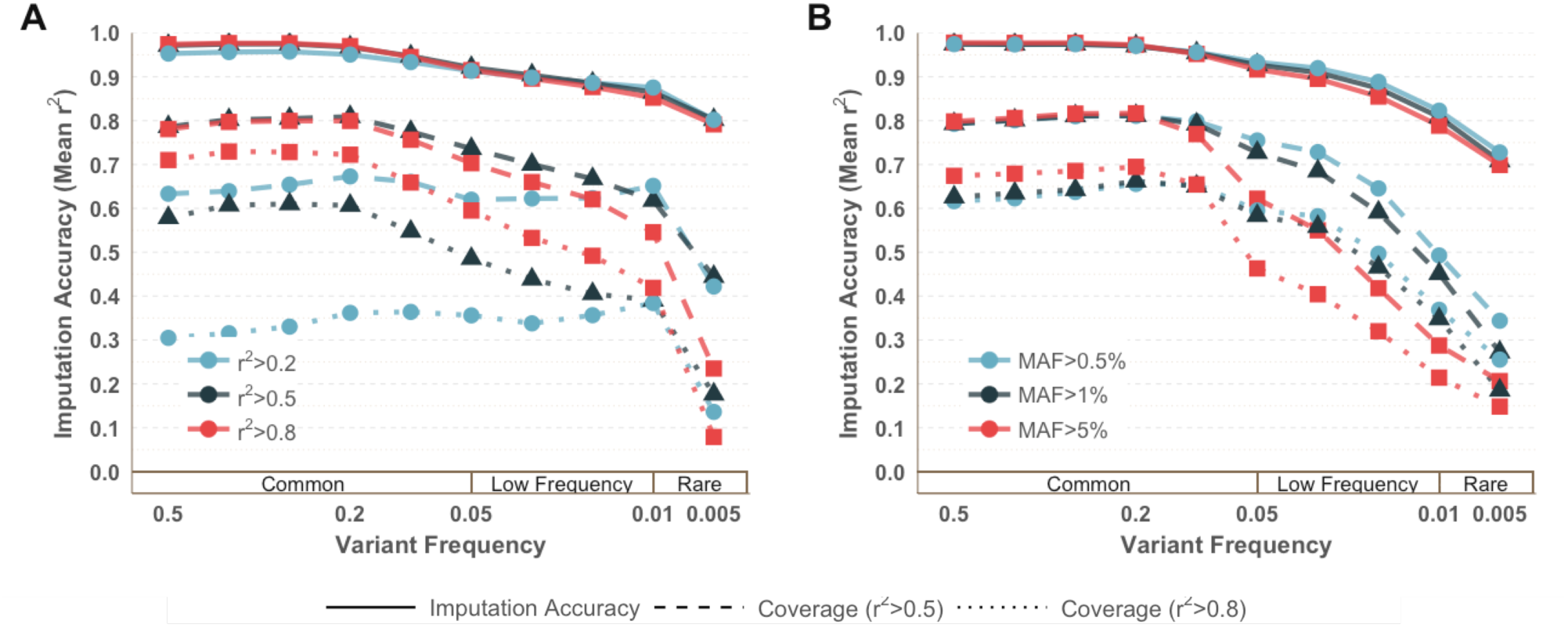
Influence of (A) minimum r^2^ threshold and (B) lower MAF threshold on imputation accuracy and coverage(r^2^>0.5 and r^2^>0.8) within populations from the Americas with an allocation of 1M sites.

The impact of minimum minor allele frequency threshold was negligible across variants with MAF>5% for all non-African populations **(Supplementary Figure 6)**. Within populations of African descent, limiting tags to variants with MAF>5% resulted in increased imputation accuracy for all frequency bins, especially for common variants. Lowering the MAF to 0.5% reduced accuracy in African-descent populations across all frequency bins. For EUR, SAS, and AMR, tags with MAF>1% had decreased accuracy for variants with MAF 0.5-1% compared to when tags are limited to MAF>0.5%. **(Figure 4B)** The lowest limit of MAF (0.5%) showed increased accuracy for rare variation but at a slight cost to the accuracy for common sites (MAF>5%). We concluded that the best balance for tag SNP selection across all populations among these was MAF>1% within the population being tagged, as the imputation accuracy was best for MAF>5% for half of the groups (AAC, AFR, EAS) and best for MAF>0.5% for the other half (AMR, EUR, SAS). **(Supplementary Table 2)** However, the overall differences in imputation accuracy was minimal, with less than 1% between all lower MAF thresholds across all sites. Again, we observed large differences in pairwise coverage, despite negligible differences when performance is evaluated by imputation accuracy. **(Supplementary Figure 6)** This is particularly striking for African-descent populations (ASW and AFR), where there were large gains of pairwise coverage for MAF>1%, compared to MAF>0.5% and MAF>5%. As previously described, African populations have shorter LD blocks and a greater absolute number of polymorphic variants compared to other populations. ^17^ Therefore, pairwise coverage underestimates performance compared to imputation accuracy, as addressed below.

### Tagging potential differs between populations

Efficient tag SNP selection is an opportunity to boost power in downstream analyses. African and out-of-Africa populations exhibit distinct genetic architecture which resulted in different performance trends. There is therefore a need to balance the representation from all global populations. However, even when cross-population performance was prioritized, it did not guarantee equal representation of all population groups within the tag SNP set. To determine the contribution of each population, we again focused on chromosome 9 (42,215 tags), equivalent to one million sites genome-wide, selected with our novel cross-population prioritization scheme. This tag SNP allocation resulted in including all tags that were informative in at least 3 to all 6 populations in the scaffold. Out of all tags for chromosome 9, 17.96% were informative in all 6 populations. **(Supplementary Table 3)** No tags were included that were informative in only one or two populations. Of tags that were informative in 5 out of the 6 super-populations, only 54% were in LD with any target sites within EAS populations, while 93% were informative in AAC populations.**(Figure 5A)** This trend is consistent with cross-population tags tending to be less informative in EAS populations compared to the other populations. When tags are informative in 3 out of 6 groups, only 18% were informative in EAS, while 75% were informative in AAC. Tags informative in only 2 of the 6 groups were likely informative in AAC and AFR, the African descent populations, while very few of them were informative for non-African descent groups, consistent with capturing differential LD patterns in African populations.^43^ When tags are stratified by MAF (0.5-1%, 1-5%, and >5%), these trends are exaggerated in the rare and very rare MAF bins. **(Supplementary Figure 7)** As expected, the very rare variation (0.5-1% MAF) was highly population-specific with no sites in this frequency bin being informative across all populations, or even 5 out of the 6 populations. ^27^ For rare variation (1-5%), tags were the least informative within EAS, with only 36% of the tags informative in 5 out of 6 populations.

**Figure 5:**
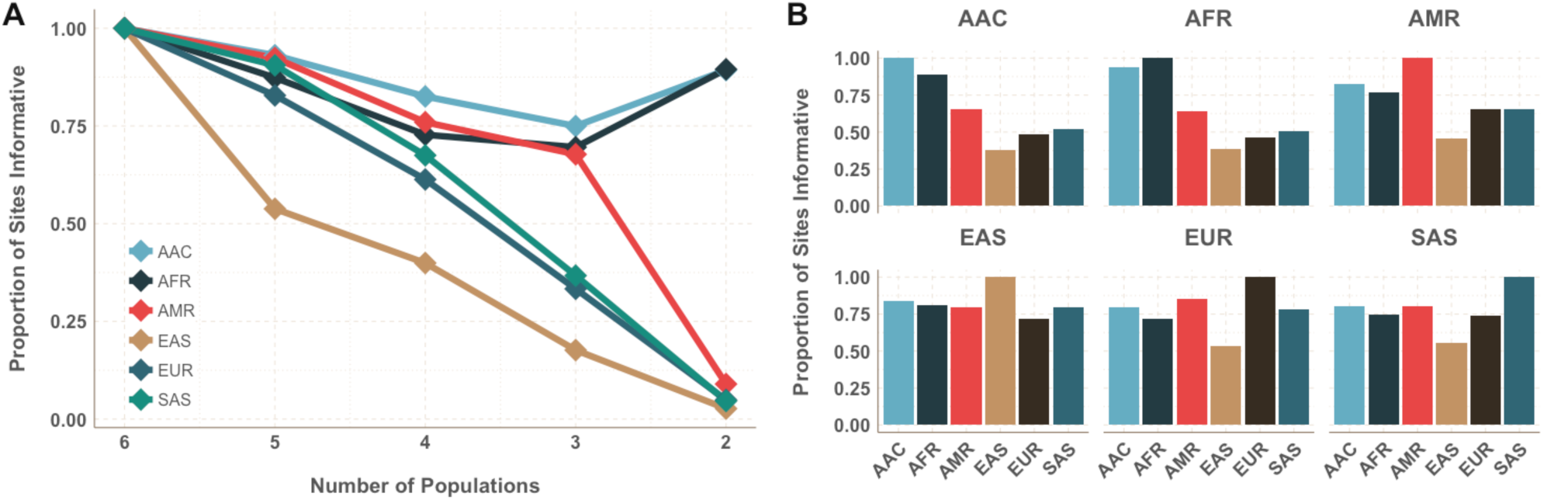
Tag SNPs informativeness across population. (A) Proportion of sites informative (r^2^>0.5, MAF>0.01, 1M site scaffold) across a number of populations, with lines corresponding to the index population. (B) Proportion of sites shared across populations, conditional on index population.

Conditional performance, or the ability of a tag which is informative in the index population also being informative in an additional population, was also examined and found to be consistent with known population histories. Of tags that are informative within AFR, 94% were informative within AAC, while only 38% were informative within EAS. **(Figure 5B)** However, among tags that were informative within EAS, 81% were informative within African populations. Once again, the stratified analyses show exaggerated trends for the very rare and rare MAF bins. **(Supplementary Figure 8)** For the very rare variation (0.5-1%), only a very small percentage (<10%) of tags are informative in other populations (AMR, EAS, EUR, SAS) if they were informative within African-descent populations (AFR and AAC). The high level of sharing between AFR and AAC is expected due to the high proportion of African ancestry within African-American and Afro-Caribbean populations. Of tags informative within EUR, 78% are also informative within AMR, largely due to the high proportion of European ancestry within some Hispanic/Latino populations. ^39,44,45^

The tags were also not equally informative in each population when it comes to the number of sites they tag with r^2^>0.5. For chromosome 9, it would take 81,416 tags to capture all possible tag-able variation with an r^2^>0.5 within AFR populations, while it would take only 28,473 tags within EAS populations to saturate coverage. However, each tag within the AFR populations captures on average 7.17 other sites, whereas for EAS populations, each tag captures on average 10.27 other SNPs. When restricting the design to a million tag SNP scaffold, each tag captures on average 16.16 other SNPs within EAS populations and 12.16 other SNPs in AFR populations. **(Table 1)** This reflects the different underlying genetic architecture of these different groups.

**Table 1:** Performance per tag SNP to capture all variation possible with r^2^>0.8 on chromosome 9, as well as within a million site genome-wide scaffold allocation through cross-population prioritization.

| Population | All Possible Tags |  | One Million Tag Scaffold |  |
| --- | --- | --- | --- | --- |
|  | Number of Tags | Sites Captured per Tag | Number of Tags | Sites Captured per Tag |
| AAC | 74,255 | 8.04 | 36,336 | 12.97 |
| AFR | 81,416 | 7.17 | 34,548 | 12.16 |
| AMR | 43,065 | 9.40 | 28,691 | 12.80 |
| EAS | 28,473 | 10.27 | 16,457 | 16.16 |
| EUR | 35,027 | 9.48 | 22,111 | 13.63 |
| SAS | 37,644 | 9.28 | 23,480 | 13.33 |

### Limits of tagging and imputation

Not all of the human genome can be captured through pairwise tagging given existing reference panels. For each super population, we filtered for sites that were polymorphic (MAF>0.5%) and had no pairwise correlation (r^2^>0.2) with any other site within one megabase. The number of these “lone sites” without any pairwise correlation was dependent upon population. AAC had the greatest number of lone sites, but that is likely due to the significantly decreased sample size compared to the other populations. **(Table 2)** The lowest number of lone sites was found within AMR. Although these sites have no notable pairwise correlation with any other site in the human genome, haplotypes may be informative and allow the recovery of information for imputation. We again assumed a one million genome-wide tag SNP scaffold allocation with minimum MAF of 1% and minimum r^2^ threshold of 0.5 and imputed to the entire 1000 Genomes reference panel. As expected, imputation accuracy and ability to recover information was population-specific. The imputation accuracy within AAC was an outlier when compared to other populations, with 80.72% of lone sites being imputed with at least the accuracy of r_acc_^2^≥0.5 and over 50% of sites being imputed with even higher accuracy (r_acc_^2^≥0.8). Many of these lone sites within AAC were captured with pairwise and haplotype LD within other populations, primarily AFR and to a lesser extent EUR. While there were likely insufficient allele counts for accurate correlation estimation within AAC due to the small sample size, this information could be recovered using a global reference panel. The number of unrecoverable “dark sites”, which had no pairwise correlation and were not recoverable with imputation using haplotype information, was the largest in EAS and is consistent with known demography and population history yielding an excess of highly rare variation compared to other populations.^27^

**Table 2:**
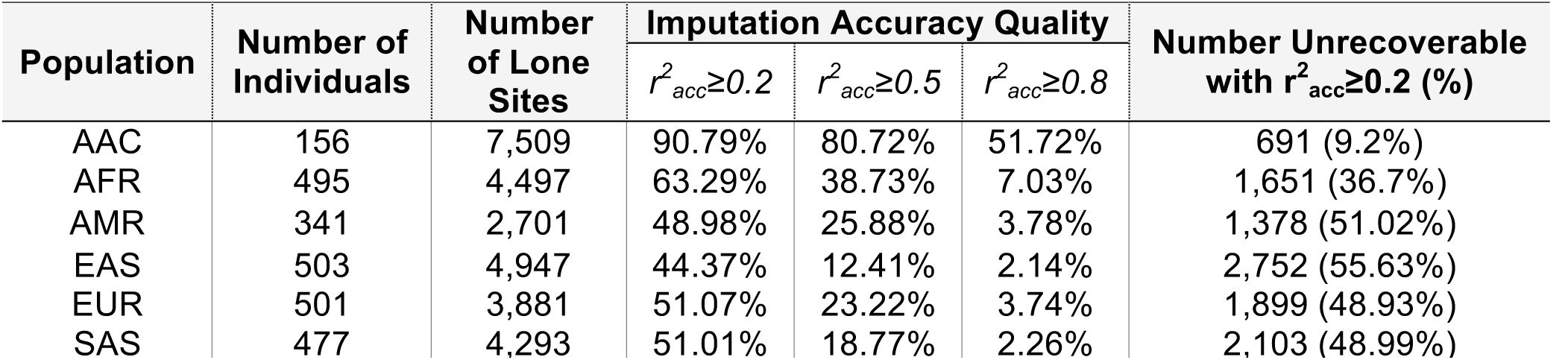
Lone sites by super population and their imputation accuracy for a 1M site scaffold.

### Pairwise coverage versus imputation accuracy

When evaluating the performance of a GWAS scaffold, there are numerous factors to take into consideration. These include the number of sites you have allocated to tag SNPs and what your priorities are for balanced representation. To a lesser extent, the benefits and pitfalls of prioritizing low-frequency variants must be weighed. However, we have demonstrated that the influence of these components are highly dependent on how performance is measured. The notion of genomic “coverage” has historically been estimated using pairwise correlations, and therefore this term will be used to denote the proportion of polymorphic sites that are in pairwise LD (r^2^ threshold) with at least one tag SNP. We calculated coverage separately per super population at an r^2^ threshold of 0.5 and 0.8 within minor allele frequency bins identical to the imputation accuracy estimation analyses, assuming a genome-wide tag SNP set of 500,000 and 1,000,000. **(Table 3)** For a tag SNP set of one million sites, coverage was lowest in AFR with an overall average of 59.15% for all sites with MAF>0.5% and r^2^>0.5. **(Supplementary Figure 9)** When the r^2^ threshold is raised to 0.8, the proportion of sites in linkage disequilibrium with at least one tag SNP lowers to 28%. **(Figure 6)** The highest coverage was found in populations from the Americas (AMR) and East Asia (EAS). For a lower r^2^ threshold of 0.5, 79.9% of AMR sites with MAF>0.5% were covered. When using the higher r^2^ threshold of 0.8, East Asian populations had the highest coverage with 63.08% of sites in LD with at least one tag SNP. This difference is even more marked when looking at a smaller tag SNP set of 500,000 sites. **(Supplementary Figure 10-11)** African populations now have an overall coverage of 33.17% with r^2^>0.5 and 14.10% with r^2^>0.8. East Asian populations have the highest coverage with 73.16% of sites covered with r^2^>0.5 and 55.09% with r^2^>0.8.

**Figure 6:**
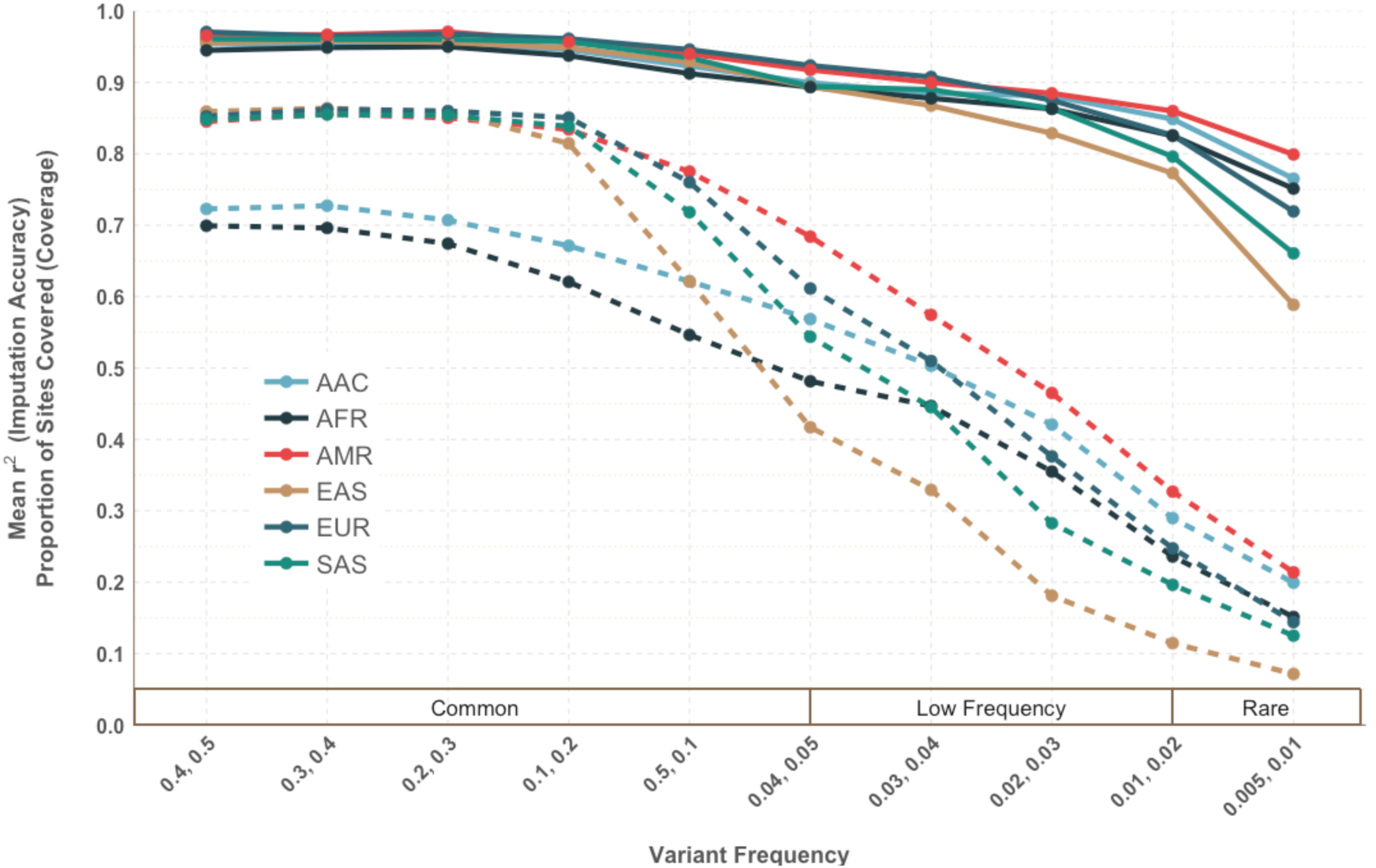
Coverage versus Imputation Accuracy, assuming a genome-wide scaffold size of one million tags. Coverage is shown with an r^2^>0.8. While pairwise tagging values are low, particularly in African-descent populations, multi-marker imputation accuracy remains high across groups.

**Table 3:** Coverage of 1 million and 500,000 tag SNP set by super population for all polymorphic sites on chromosome 9 with MAF>0.5%

| Super population | Total Number of Polymorphic Sites | Scaffold of 1,000,000 tags |  |  | Scaffold of 500,000 tags |  |  |
| --- | --- | --- | --- | --- | --- | --- | --- |
|  |  | Coverage |  | Imputation Accuracy | Coverage |  | Imputation Accuracy |
| | | $r^2>0.5$ | $r^2>0.8$ | | $r^2>0.5$ | $r^2>0.8$ | |
| AAC | 780896 | 63.64% | 30.27% | 90.59% | 34.03% | 14.07% | 84.85% |
| AFR | 777207 | 59.15% | 28.05% | 89.62% | 33.17% | 14.10% | 83.32% |
| AMR | 503804 | 79.90% | 53.60% | 92.77% | 61.00% | 37.02% | 90.09% |
| EAS | 367189 | 76.95% | 63.08% | 86.28% | 73.16% | 55.09% | 84.16% |
| EUR | 414184 | 78.77% | 62.65% | 91.02% | 72.87% | 52.86% | 88.90% |
| SAS | 455573 | 74.84% | 56.97% | 88.09% | 67.28% | 45.91% | 85.46% |

These trends are in striking contrast to those we observed in imputation accuracy. When comparing a tag SNP set of 1 million, pairwise LD coverage is the lowest in populations of African descent (59% with r^2^>0.5) yet imputation's ability to recover un-typed sites is on average high and consistent with other populations (imputation accuracy of 89.62%) among SNPs with a minor allele frequency above 0.5%. This contrast is also found in East Asian populations, which had one of the highest proportion of polymorphic SNPs with r^2^>0.5 for coverage (76.95%), but the lowest imputation accuracy (86.28%). **(Table 3)** When sites are stratified by minor allele frequency bins, the differences in trends are even more striking. **(Figure 6, Supplemental Figure 9)** For example, within the lowest frequency bin (0.5% to 1%) for admixed populations of African-descent, the coverage of sites for a set of 500,000 tag SNPs with r^2^>0.8 falls below 10%, however the imputation accuracy remains relatively high at 77.82%. These trends are consistent and more dramatic when evaluated within a tag SNP set of 500,000 sites. **(Supplemental Figures 10-11)** These observations reinforce the necessity of examining imputation accuracy, instead of pairwise coverage, when evaluating the performance of tag SNPs.

## Discussion

As larger and larger whole genome reference panels come into availability for imputation, it is important to design arrays with this ultimate goal in mind. There are currently two accepted methods of evaluating the performance of a tag SNPs: pairwise LD “coverage” and imputation accuracy. Coverage has historically been used as a term to denote the proportion of polymorphic sites that are in linkage disequilibrium with at least one tag marker above a certain r^2^ threshold. ^46–49^ Genotyping arrays are typically compared using this score averaged across the genome. However, as others and we have demonstrated, restricting performance assessment to this definition of pairwise coverage is limited by removing multimarker information. ^33,34^ Evaluating imputation accuracy, particularly via leave-one-out cross validation, is highly computationally intensive, but provides a better assessment of how well untyped variation can be recaptured and a more realistic depiction of array performance. Imputation accuracy is also a more useful statistic in a practical sense, especially with the development of deeper and more diverse reference panels,^17,18,50–52^ as performing GWAS with imputed variants is now the expectation. Emerging evidence suggests that very rare variants that are poorly tagged by an individual tag SNP will be accessible via imputation, due to added haplotype information, particularly as sample sizes move beyond the thousands into the tens of thousands. ^19,34^.

In addition, previous tagging strategies have predominantly focused on optimizing performance in a single population. In prioritizing potential tags by their ability to provide linkage disequilibrium information across multiple populations, we were able to demonstrate that cross population tag SNP selection outperforms single population selection. This boost in imputation accuracy exists across all populations and frequency bins. We simulated tag SNP sets for a range of sizes (250,000-2 million), as well as for several minimum minor allele frequencies (0.5%, 1%, 5%) and minimum r^2^ thresholds (0.2, 0.5, 0.8). For investigators with limited real estate or budget for tag SNP selection, we found that the biggest improvement in imputation accuracy provided with our cross population approach was with the smaller array sizes (250K) when compared to a naïve design or biased population ascertainment. As expected, the influence of MAF and r^2^ threshold was population-specific. For African-descent populations, including tag SNPs with a low threshold of r^2^ ≥ 0.2 resulted in lower imputation accuracy across all bins, while in other populations (EUR, AMR, SAS) tags at r^2^ ≥ 0.2 led to increased imputation accuracy for low frequency variants to the detriment of common variation. This is due to the lower LD patterns overall in African haplotypes, requiring denser coverage. The best balance was found with a moderate r^2^ threshold of ≥ 0.5 for those seeking to perform well across all populations. This compromise is also present in choosing the lower MAF threshold. Limiting tag SNP selection to common variants with MAF ≥ 5% produced the highest imputation accuracy across all frequency bins within African-descent populations. However, this threshold decreased imputation accuracy for low frequency and rare variants in all other populations. Therefore, the best balance is once again found in the moderate value of MAF ≥ 1%. Investigators will need to take their priorities into account when selecting the correct thresholds for their populations and if they have a specific target frequency bin. We chose to prioritize all populations equally to provide a design of broad global utility, but if the study is comprised of mostly one ancestral group then the investigators should choose the appropriate thresholds tailored for their study.

Consistent with demographic history, the potential to capture variation with a limited allocation is unequal between the different populations in the 1000 Genomes Project. The naïve tagging approach will bias tag SNP selection to be primarily informative within African-descent populations. The absolute number of polymorphic sites within African populations is much larger than other populations, and while LD tends to be lower than in other populations, the high number of potential tags and pairwise correlations overwhelms the other populations’ contributions without controlling for this unique pattern. By prioritizing potential tags that provide information across all populations, the population-level contributions are more balanced without detriment to the African-descent groups (**Figure 4**). The absolute number of rare variants (MAF < 1%) is larger in African populations, but the frequency spectrum is more skewed towards rare variants in populations with recent bottlenecks and exponential population expansion, such as in East Asians. Contrasting these two populations (AFR and EAS), East Asian populations require fewer sites to saturate coverage, with each potential tag being in LD with more sites. However, far more polymorphic sites across the genome cannot be captured with either pairwise linkage disequilibrium or through haplotype information with imputation accuracy within these populations due to a dearth of LD information. This is amplified by the lack of comprehensive reference panels for many populations, such as East and South Asia. As reference panels are expanded, more variation will be captured to inform tag SNP selection and imputation accuracy, and we expect imputation accuracy to improve for all populations and across the frequency spectrum.^19^

As cosmopolitan biobanks and large-scale multi-ethnic epidemiological studies become more commonplace, the available technology to capture genetic variation must keep pace. It is important to rely on a platform that is as equitable as possible in providing information about the groups of interest when conducting genetic association studies within diverse populations. There are various considerations that an investigator must consider when selecting tag SNPs to customize or build an array, or evaluate an available commercial array. Assessing performance through imputation accuracy, as performed here, is the most apt comparison between tag SNP sets, allowing a real-world look at the extent of variation a set of tags can capture based on haplotype structure. We have presented an improved tagging algorithm and evaluation pipeline that prioritizes cross-population performance for increased imputation accuracy across multiple populations and the full range of MAF ≥ 0.5%. We also provide recommendations and context for other researchers interested in similar goals.

The power to identify relevant loci is inherently constrained by sample size and genome coverage. Imputation improves this by providing increased effective coverage across the genome. It is important to note that algorithmic development both on association testing and imputation methods has been a productive avenue of research since GWAS began, with new methods providing incremental improvements in statistical power. Here, we demonstrate with a fixed allocation, methods to improve statistical power by tailored SNP selection in the initial array, with sometimes dramatic improvements in imputation accuracy. With the expansion and improvement of global reference panels, genotyping arrays will be able to capture an increased amount of variation, especially when cross-population performance is prioritized. The unified framework presented will enable investigators to make informed decisions in the development and selection of genotyping arrays for future large-scale multi-ethnic epidemiological studies. This increased representation of multi-ethnic genetic variation will promote the investigation of the genetics of complex disease and the improvement of global health in the next phase of GWAS.

## Author Contributions

GLW, CF, RKW, CC, GA, HMK, MB, CDB, CRG, and EEK conceived of and designed the experiments. GLW performed the data analysis. CR, DT, RW, ARM, SS, and HMK contributed analysis tools/materials. GLW wrote the manuscript with comments from CF, ARM, LH, CC, GA, CRG, and EEK. All authors read and approved the final manuscript.

## Acknowledgments

Research reported in this paper was supported by the Office of Research Infrastructure under award number S10OD018522 and the National Human Genome Research Institute under award numbers U01HG007376, U01HG007417, and R01HG000376 of the National Institutes of Health. CRG was supported partially by T32HG00044. The content is solely the responsibility of the authors and does not necessarily represent the official views of the National Institutes of Health.

## Materials and Methods

### Genetic Data

The genetic data are from the 1000 Genomes Project (1000 Genomes) Phase 3 data release, version 2 (7/8/2014) containing whole genome sequences for 2,535 individuals from 26 global populations. ^53^ Sequence data were in VCFv4.1 format, mapped to the forward strand and variants annotated as reference or alternate alleles. Only biallelic SNPs were included in this analysis (77,224,748 SNPs total). A list of known cryptically related individuals was obtained from the 1000 Genomes FTP site, and one individual from each related pair were subsequently removed (n=62). Individuals were assigned to their super populations according to the original 1000 Genomes assignments (EAS=East Asian, EUR=European, AFR=African, SAS=South Asian, AMR=Americas, comprising 503, 501, 495, 477, and 341 individuals, respectively). Two populations of admixed African ancestry (ASW and ACB) were removed from the African super population and formed a separate African American/Caribbean (AAC) super population (n=156).

### Tag SNP Selection

Allele frequency was estimated within super population for each SNP using Plink v1.9. ^54^ Linkage Disequilibrium (LD) was also calculated within each super population using Plink v1.9 and settings for pairwise linkage with a minimum r^2^ of 0.2 within a maximum distance of 1 megabase (mb). Tag SNP selection was performed per chromosome in the program TagIT ^23^, with frequency and LD files for each super population as input. The TagIT algorithm analyzed each super population separately. After filtering based on the minor allele frequency (set as either 0.5%, 1% or 5%), TagIT annotates the tag SNP that has the highest number of LD pairs with r2 above a minimum threshold (set as either 0.2, 0.5, or 0.8). The selected tag SNP and all of its linked SNPs are masked and TagIT finds the next tag SNP with the highest number of LD pairs. The output for each super population included for each index tag SNP the number of sites in LD, as well as the number of unique sites that weren't already tagged by a previously chosen tag. The number of unique SNPs tagged across all populations per tag SNP was tallied in the final output.

### Cross-population tag SNP ranking and scoring

The naive approach ranked potential tags by the absolute number of unique SNPs that are tagged across all super populations. From this list, the top SNPs were selected for the appropriate allocation. To ensure performance of the tags across multiple populations, the cross-population prioritization schema first ranks tags by the number of populations in which they are informative, meaning they tag at least one site (Supplementary Figure 1). This ensures that the top ranked SNPs are not biased to a super population with large LD blocks or high SNP density in which one tag can contribute information about many other SNPs. Within each one of these categories (all 6 super populations down to only 1 super population), the tags are ranked by the number of unique tags across all six super populations, as was done in the original approach. The appropriate allocation is selected from the top of this list, scaled to the size of the chromosome of interest.

### Metric of Performance

Coverage and imputation accuracy were assessed using all polymorphic biallelic sites within the 1000 Genomes Phase 3 data release, version 2. Sites were categorized into ten discrete minor allele frequency bins: (0.005-0.01], (0.01-0.02], (0.03-0.04], (0.04-0.05], (0.05-0.1], (0.1-0.2], (0.2-0.3], (0.3-0.4], and (0.4-0.5]. The term "coverage" is used to denote the proportion of untyped sites that had at least one tag SNP with pairwise r^2^ greater than a certain threshold (0.2, 0.5, or 0.8). Imputation accuracy was determined through a leave-one-out internal validation approach with the 1000 Genomes Project Phase 3 data using a modified version of Minimac.^19^ Correlation was calculated comparing the estimated dosages to the true genotypes from the original vcf files.

### Ascertainment Bias Analyses

Population-specific tags were selected separately through TagIT for each super population with a genome-wide allocation of 500,000 sites. All tags had a minimum MAF of 1% and a minimum r^2^ threshold of 0.5. Each of the single population ascertained tag lists assessed for imputation accuracy in all six super populations, including their index population. Imputation accuracy was calculated as previously described and limited to chromosome 9.

### Local Ancestry

Local ancestry was estimated using RFMix ^55^ assuming three ancestral backgrounds: African, European, and Native American, and is described in detail in ^42^ Tracts were dropped if smaller than 20 cM to improve accuracy in local ancestry estimation. Diploid ancestry with three ancestral backgrounds yielded six categories of variation. Imputation accuracy was then calculated separately per diploid tract category, with all other sections masked out. Results were aggregated across all chromosomes to calculate the genome-wide performance per diploid ancestry. Tracts were removed from analysis if the ancestral diplotype was found in fewer than 5 individuals. This included AFR-NAT and EUR-NAT within ACB which only occurred in 2 individuals each, NAT-NAT diplotypes in ASW which occurred in one individual, and AFR-AFR diplotypes in MXL which occurred in 3 individuals.

### Cross-population patterns of linkage disequilibrium

To determine how many sites were in LD with tag SNPs across all 6 super populations, we selected one million SNPs for a GWAS scaffold using a minimum r^2^ of 0.5 and a minimum MAF of 0.01 on chromosome 9. We calculated the number of polymorphic sites (MAF>0.5%) and the proportion of these sites that were in LD (r^2^>0.5 or r^2^>0.8) with at least one tag marker. To determine sharing of tags across multiple populations, we calculated the proportion of tag markers that were informative in other populations, conditional upon them being informative in the index population. The proportion of sites shared among multiple populations was calculated as the proportion of tag SNPs that performed in a certain number of populations (from 1 to 6 super populations) per super population.

### Tagging Potential

Tag SNPs were selected with a minimum r^2^ of 0.5 and a minimum MAF of 0.01 on chromosome 9. The potential for tagging was determined assuming an infinite site scaffold, using all possible tags until every pairwise relationship with r^2^ above 0.5 was captured. The average number of sites captured per tag was calculated in each super population separately, using only the tags that were informative within that population. We also calculated these trends assuming a scaffold of one million sites, following the same procedures. The “dark sites” were calculated as sites in which there was no pairwise correlation with any other site with r^2^>0.2, determined separately for each super population.

